# Evoked and spontaneous pain assessment during tooth pulp injury

**DOI:** 10.1101/742049

**Authors:** Heather Lynn Rossi, Lily Pachanin See, William Foster, Saumitra Pitake, Jennifer Gibbs, Brian Schmidt, Claire H. Mitchell, Ishmail Abdus-Saboor

## Abstract

Injury of the tooth pulp is excruciatingly painful and yet the receptors and neural circuit mechanisms that transmit this form of pain remain poorly defined in both the clinic and preclinical rodent models. Easily quantifiable behavioral assessment in the rodent orofacial area remains a major bottleneck in uncovering molecular mechanisms that govern inflammatory pain in the tooth. Here we use a dental pulp injury model in the mouse and expose the tooth pulp to the outside environment, a procedure we have previously shown produces pulpal inflammation. We demonstrate here with RNAscope technology in the trigeminal ganglion of injured mice, an upregulation of genes that contribute to the inflammatory pain state. Using both evoked and spontaneous measures of pain in the orofacial area, including application of von Frey Hair filaments and pain feature detection with the mouse grimace scale, we reveal a differential timeline of induction of spontaneous pain versus mechanical allodynia following pulpal injury. This work demonstrates that tooth pain can be easily assessed in freely behaving mice using approaches common for other types of pain assessment. Harnessing these assays in the orofacial area during gene manipulation should assist in uncovering mechanisms for tooth pulp inflammation and other forms of trigeminal pain.

## Introduction

Pain from the infected tooth pulp (pulpitis) can be unrelenting and many patients report this form of pain as the most intense type of pain they have ever experienced^1^. Mechanical hypersensitivity of the tooth is associated with greater pain intensity ratings overall ^2^. Prevailing treatment options for painful pulpitis consists of pulp or tooth removal, which can have lasting consequences for dental function and in some patients there may still be lingering pain ^3,4^. Therefore, there is a critical need for development of new therapeutic approaches that alleviate tooth pain while leaving pulpal issue intact and avoiding complex dental procedures. Moreover, untreated ongoing inflammation of the pulp can lead to more widespread nociceptive hypersensitivity in trigeminal tissues, an issue further compounded in individuals who cannot afford proper dental treatment^5^. The era of new innovative approaches to treat tooth pain will be driven by an increase in our fundamental understanding of the genes and neural circuit pathways that drive tooth pain states. However, we first need to establish feasible and objective behavioral paradigms that measure pain in the orofacial area in preclinical rodent models.

Clinically, mechanical hypersensitivity and spontaneous pain are particularly problematic for patients ^2-4^, and there are behavioral assessment tools for these in rodent models. To date, only a handful of studies using the tooth pulp injury model have examined mouse behavior, and these studies have not incorporated some of the common assays used to measure pain hypersensitivity ^6-8^. The predominant assessment tool for mechanical pain measurement in rodents are reflexive withdrawal assays in which calibrated von Frey hair filaments (VFHs) are applied to the hind paw and the experimenter decides which filaments evoke an animal’s withdrawal ^9^. We have recently improved on this method by incorporating high-speed videography, statistical modeling, and machine learning to more objectively assess the mouse pain state following hind paw stimulation^10^. VFHs can also be applied to the face, but this presents more challenges because the animal’s attention is more engaged with the stimulus, as we have previously experienced ^11^. However, recent elegant work in freely behaving mice used both VFH stimulation of the whisker pad and optogenetic activation of trigeminal nociceptors to uncover a craniofacial neural circuit for pain^12^, so it is not impossible.

Another approach to measure spontaneous pain in rodents is a paradigm called the Mouse Grimace Scale (MGS), that interrogates facial expressions including the positioning of the mouse nose, cheek, ear, eye, and whiskers^13,14^. An advantage of the MGS over reflexive assays is that spontaneous pain resembles pain reports in the clinic and facial expressions are used in the clinic to measure pain in infants, although these assays are currently not as high-throughput in rodents as delivering stimuli and recording immediate responses. Together however, both reflexive and spontaneous measurements of pain provide advantages in that they can be performed without anesthesia, invasive implants, or time intensive tasks performed by the animal that require long term learning and memory, which may mean that interpretation of the behavior may be confounded by factors outside of the animal’s pain level.

Here we take advantage of existing pain assays, with some custom modifications, and adapt them for behavioral analyses during tooth pulp inflammation. After morphologically confirming our tooth pulp injury model, we used RNAscope technology to determine the time course of changes in molecular mediators of nociception relative to behavioral changes. To the best of our knowledge, this is the first study using the MGS to evaluate pain following dental injury, and our results revealed the occurrence of spontaneous pain within the first day following dental injury. We also adapted previous facial Von Frey methods^15^ to evaluate mechanical sensitivity, relying on the published scoring scheme, as well as the animals’ willingness to put its head through a custom designed chamber with an adjustable opening for stimulation. Because the mice can decide if they want to expose their faces to the stimuli, we were also able to record the threshold in which mice are no longer inquisitive enough to tolerate facial stimuli, and this pain threshold was able to segregate injured versus sham mice. Interestingly, these two assays present a different time course following injury, indicating spontaneous pain early and throughout the 6 day observation period, while mechanical allodynia is delayed. Taken together, the behavioral assays we have defined here to assess tooth pain should make it easier for researchers to adopt these approaches to aid in uncovering mechanisms for tooth pain.

## Results

### Morphological and gene expression changes twenty-four hours after tooth pulp exposure

In order to study inflammatory tooth pain, we used the dental pulp injury (DPI) model that we have previously described^6^ in which the dental pulp of one maxillary molar tooth is mechanically exposed using a dental drill, producing pulpitis. We began our first analyses 24 hours post-injury, and confirmed controlled removal of enamel and dentin and exposure of the pulp occurred in the molars of DPI mice (Fig.1A,B). Foreign material was microscopically present in the tooth cavity of all 3 DPI mice, sometimes in contact with the pulp (material next to arrow in Fig.1B), demonstrating that an exposed pulp collects materials from the mouse’s outside environment. Importantly, the pulp was still present and clearly exposed, and not yet necrotic, within the injury site at 24 hours (arrow Fig.1B).

**Figure 1.**
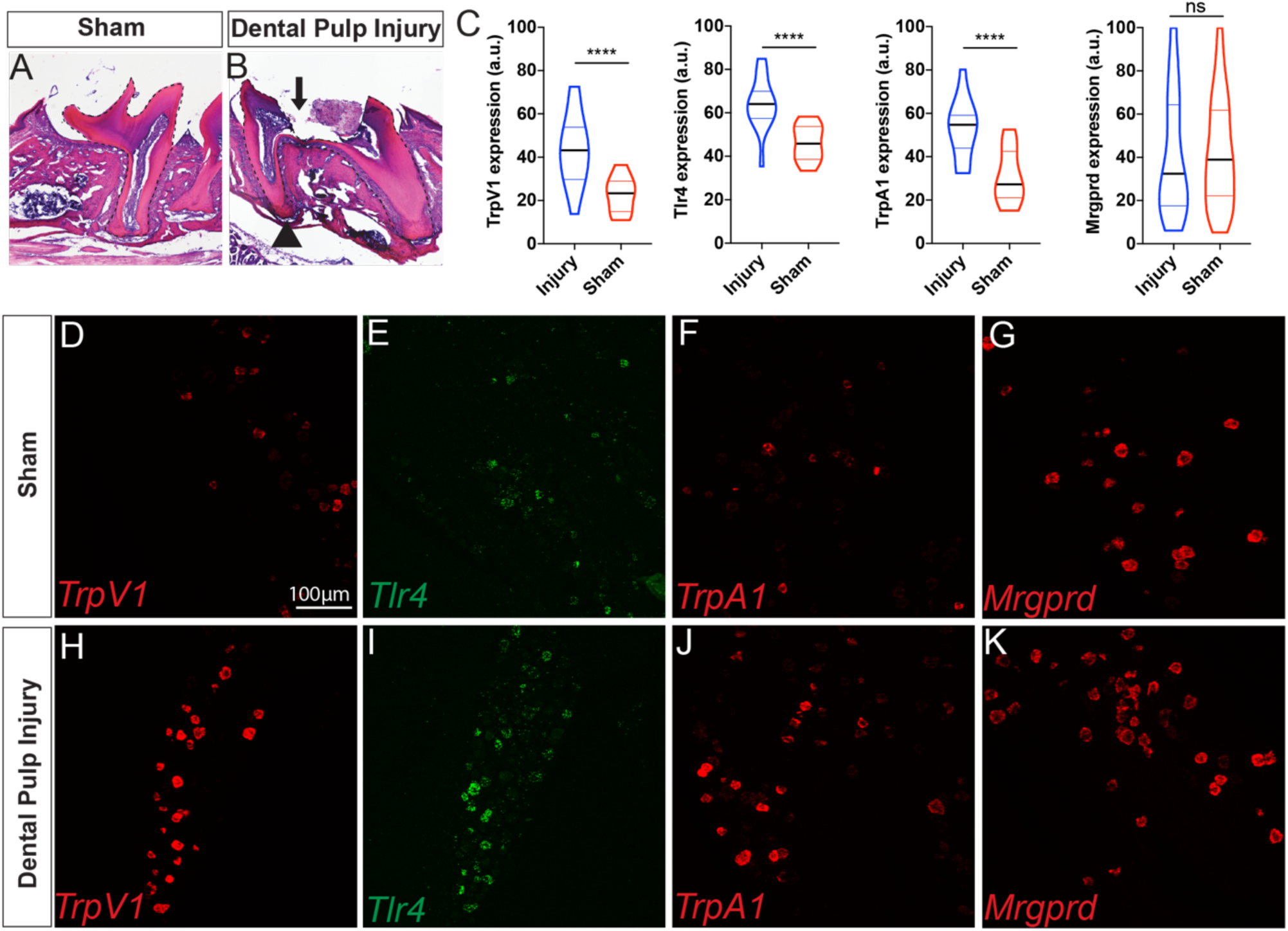
Changes in tooth morphology and trigeminal ganglia transcript levels 1 day following pulp injury. (A) Intact first maxillary molar from a sham animal and (B) injured maxillary molar with pulp exposure evident. Dotted outline marks the molar and the arrow indicates the exposure site. Healthy pulp is still evident on either side of the opening, and compacted foreign material (next to arrow) was present in the cavity. (C) Violin plot of quantified fluorescence intensity measured in arbitrary units (a.u.) defined in ImageJ for TrpV1, Tlr4, TrpA1, and Mrgprd in the trigeminal ganglia of injured (blue, n =3 mice) or sham/naive (red, n = 3 mice) mice. (D-G) Representative images of RNAscope in situ hybridization following sham and (H-K) 1 day after dental pulp injury. **** is p<0.0001 for an unpaired T-test.

Next, we utilized RNAscope technology for a sensitive read-out of RNA levels in the trigeminal ganglion of genes implicated in both nociceptive and inflammatory responses. The cell bodies of the primary afferent neurons that innervate the dental pulp reside in the trigeminal ganglion^24^. We chose to assess the Toll-like Receptor 4 (Tlr4), transient receptor potential channels vanilloid 1 and ankyrin 1 (Trpv1 and Trpa1), and the mas-related G protein coupled receptor D (Mrgprd), because all are found in neurons that innervate the dental pulp ^7,20,22,25-27^ and could be involved in the development of either spontaneous or mechanical pain in the context of infection and injury. In particular, TLR4 is part of a larger class of receptors that recognize pathogen- and damage-associated molecular patterns (PAMPs and DAMPs) ^19^, and has known interactions with both TRPV1 and TRPA1 in the context of dental injury ^20-23^. A direct role for Mrgprd in dental injury-related pain has not been established, but is possible given its expression in dental afferents ^25^ and its role in cutaneous mechanical pain perception ^28^. We found that the PAMP/DAMP family member Tlr4 was upregulated in DPI versus sham mice (Fig. 1C,E,I), as were the associated nociceptive channels TrpV1 (Fig. 1C,D,H), and TrpA1 (Fig. 1C,F,J). However, we did not observe an increase in the mechanosensitive nociceptor marker Mrgprd at 24-hours following DPI, suggesting that gene expression changes of this nociceptive membrane protein may not be driving the earliest phases of pain in the DPI model (Fig. 1C,G,K).

### Mouse grimace scale reveals presence of spontaneous pain beginning one day following pulp exposure

To assess spontaneous pain in freely behaving DPI and sham mice, we moved mice into clear custom-made chambers and recorded video of their faces. Still images were selected from these videos for assessment with the MGS (Fig. 2A). All mice, regardless of treatment exhibited very low MGS scores at baseline, which was not different between the assigned treatment groups and was not significantly affected by the sham treatment (Fig.2). We found a significant increase in the MGS at all post-exposure time points captured (Fig.2C, there was a significant effect of time, F_3,30_ = 5.776 p=0.0031, treatment F_1,10_ = 18 p=0.0017, and a significant interaction, F_3,30_ = 12.75 p<0.0001). The MGS features that differed in the DPI group versus sham were the pulling back of the ears, nose and cheek bulging, as well as orbital tightening (Fig. 2B). These results are consistent with the previous report showing that MGS scores are highest for pain emanating from internal organs ^13^, demonstrating that this assessment tool can be successfully co-opted for painful pulpitis. This finding also indicates that mice experience ongoing pain within the first day following pulp exposure that persists throughout the observation period.

**Figure 2.**
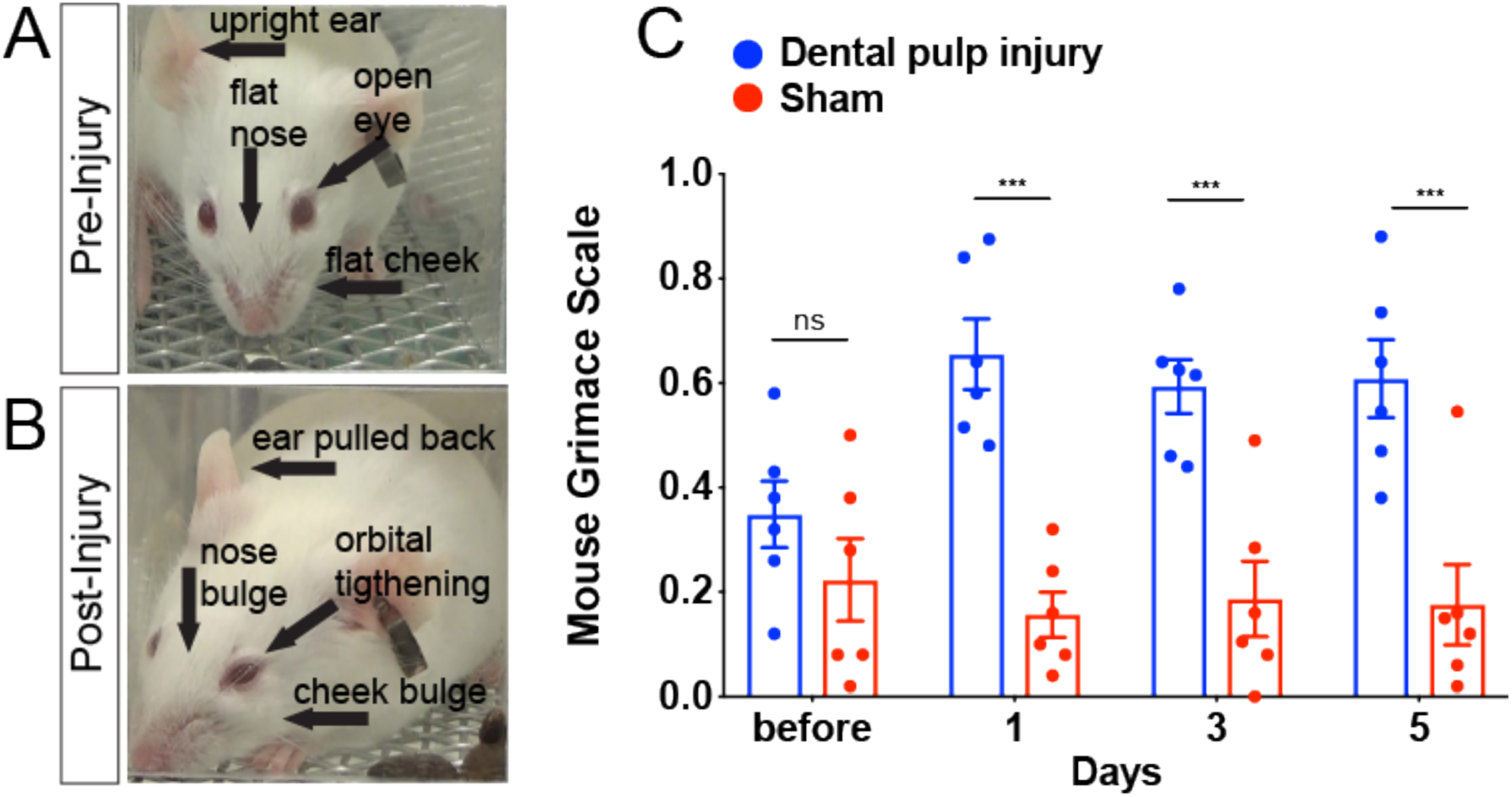
Mouse Grimace Scale following pulp exposure. (A) Before injury mice have low or no score for each of the action units, while after injury (B) prominent presence of the action units, as labeled on the example images from the same mouse. (C) We found a significant increase in the Mouse Grimace Score at all post-exposure time points captured, n = 6/treatment, Dental Pulp Injury (blue) and sham (red). * indicates p<0.0001 within the pulp exposed group before vs. after (Dunnett’s post-hoc).

### Mechanical allodynia in the face is fully developed by day 4 post pulp exposure and worsens

To determine how mice respond to evoked stimuli, we applied von Frey hair filaments to the skin between the whisker pad and eye in DPI and sham mice and recorded nocifensive behavioral responses associated with withdrawal from the stimulus. Although our stimulus did not touch the tooth directly, we hypothesized that we might observe hypersensitivity in the orofacial skin surrounding teeth, which would indicate a more widespread trigeminal sensitization, reminiscent of findings in the clinic when treatment complications arise^29,30^. Additionally, mice had to poke their heads through our custom-made chambers to allow the VFHs to make contact with the facial skin. Using this paradigm, we observed that unilateral exposure of tooth pulp on one molar changes response scores and thresholds for Von Frey stimulation on both sides of the face suggestive of mechanical allodynia (Figs.3-5). The earliest significant change was an increase in response scores across all VFHs averaged together, at day 2 on the contralateral side (Fig.3B, Contralateral: a significant effect of time: F_3,30_ = 6.43 p =0.0017, a significant interaction: F_3,30_ = 7.69 p=0.0006, but no significant effect of treatment alone: F_1,10_ = 2.272 p =0.1626). By day 4, response scores are significantly increased in pulp-exposed mice on both sides, which persists on day 6 (Fig. 3B, Ipsilateral: significant effect of time: F_3,30_ = 3.302 p =0.0336, no significant effect of treatment alone: F_1,10_ = 2.34 p =0.1571, but significant interaction: F_3,30_ = 4.836 p=0.0073). Overall this indicates that injured mice, but not shams, exhibit a gradual increase in response scores that is most evident on day 4 and maximal on day 6.

**Figure 3.**
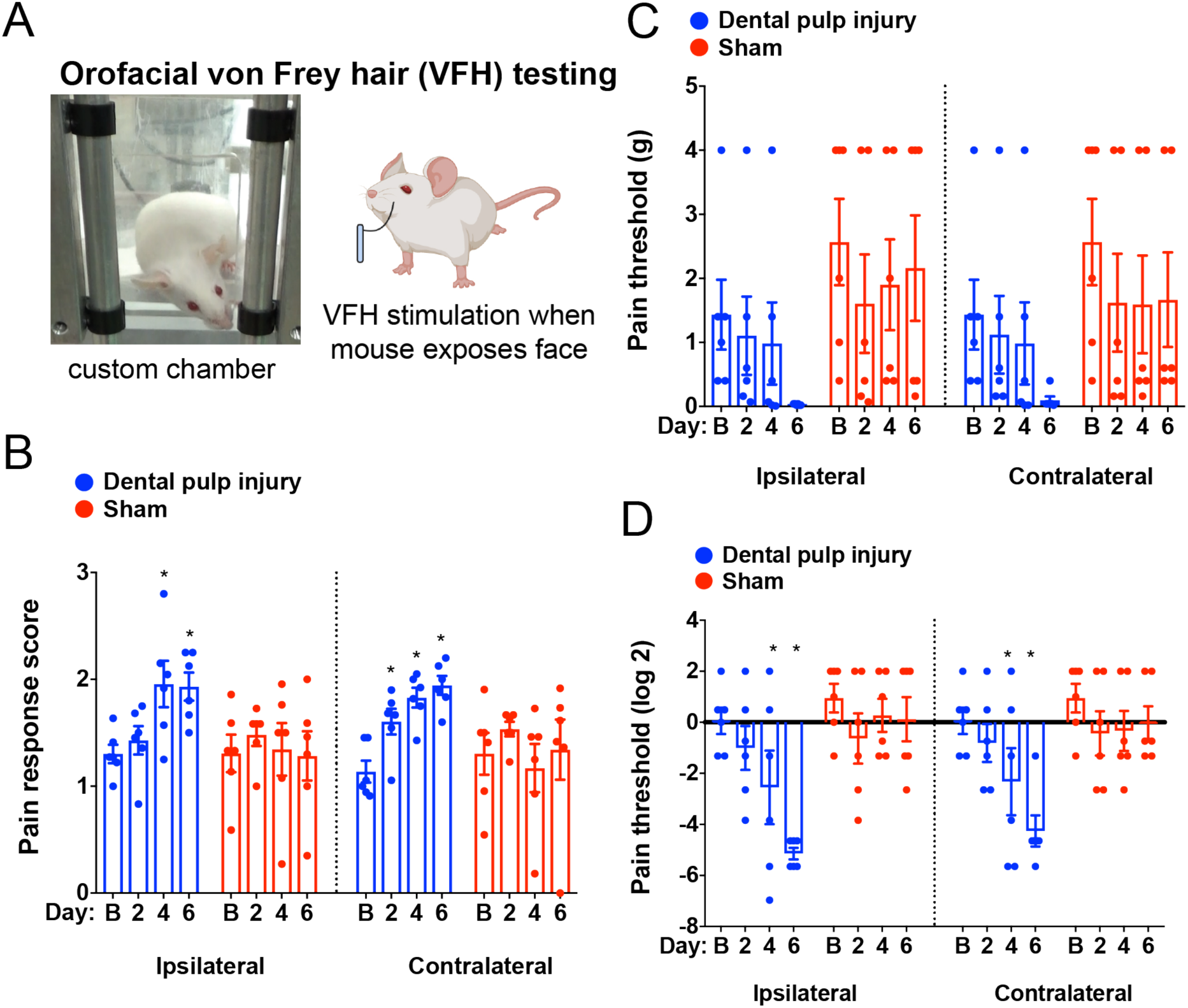
Facial Von Frey apparatus, response scores, and threshold changes following pulp exposure. (A) The mice were placed in the chamber for testing, which has adjustable openings that can be closed between tests and allows mice to put their head past the bars if they choose. When mice choose to expose their faces they are stimulated as depicted with a von Frey hair. (B) After dental pulp injury (blue) there is a significant increase in pain response scores across all of the filaments tested, which is not observed in sham-treated mice (red). This is evident from day 4 onward on the ipsilateral side, and on day 2 onward on the contralateral side. (C) Raw threshold are significantly different from normality according to the Shapiro-Wilks Test, but a decrease with time is apparent in the injured group not observed in sham mice. (C) After log transformation of threshold data to conform to normality, we find a significant decrease in threshold following pulp exposure on both sides, evident from day 4 and significantly lower than shams at day 6. * indicates p<0.05 for the indicated time point versus baseline (Dunnett’s post-hoc).

We also examined the threshold where animals either scored a 3 or refused stimulation. Raw thresholds (Fig. 3C) were log-transformed to better conform to normality (Fig. 3D) and statistical analysis of transformed data indicated a significant decrease from baseline threshold on both ipsilateral and contralateral sides beginning at day 4 post pulp exposure. On day 6, the thresholds of pulp exposed mice were significantly lower than shams (a significant effect of time: ipsilateral - F_3,30_ =6.981 p =0.0011, contralateral - F_3,30_ =6.842 p=0.0012, of treatment ipsilateral - F_1,10_ =7.424 p =0.0214, contralateral - F_1,10_ =4.981 p = 0.0497, and interaction ipsilateral - F_3,30_ =5.599 p =0.0036, contralateral - F_3,30_ =4.01 p=0.0164).

Next, we analyzed our Von Frey data in two different ways guided by the response score and the “break point” built into the assay design. The apparatus allows the mouse to learn over time that their natural urge to explore may result in mechanical stimulation to a hyperalgesic area, at which point they might choose not to expose their face through the opening and no stimulation would occur, i.e. their “break point”. This may occur in the sham group also at the higher stimulus intensities as they are repeatedly tested. There is a large degree of disagreement in the field regarding what filament ranges constitute normally “painful” vs “non-painful” stimulation in the absence of injury or damage, which we have attempted to address for hind paw stimulation with VFHs ^10^. Often this determination is made arbitrarily by the investigators based on human perception. Here, we use the Von Frey response scores and the break point to determine what range of filament weights correspond to a “non-painful” versus “painful” range. First, we examined the response scores by the weight of each VFH filament, to determine if they were higher across all intensities, which would suggest the presence of allodynia and hyperalgesia. Pulp exposed mice exhibit increasing response scores over time, particularly at lower filament weights (0.008-0.16g) on both sides (Fig.4A,C), which is not exhibited by the sham mice (Fig.4B,D). This seems to indicate mechanical allodynia in the injured group.

**Figure 4.**
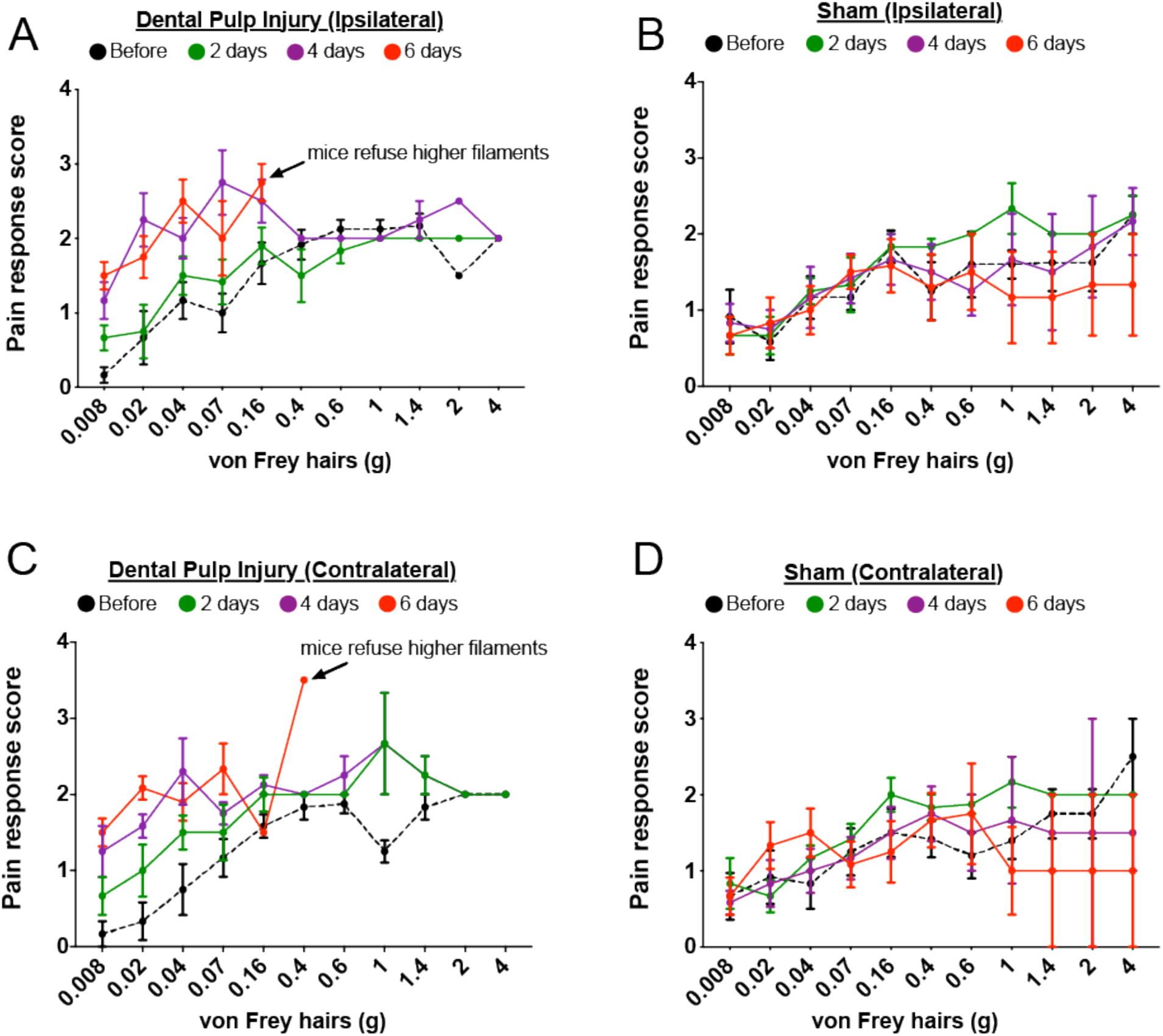
Von Frey pain response score by filament intensity and day post-injury or sham. (A) Ipsilateral and (C) contralateral response scores for low intensity filaments are increased at day 4 (purple) following injury, and by day 6 (red) mice met threshold criteria or refused filaments higher than 0.16 or 0.4g. In contrast, (B) ipsilateral and (D) contralateral scores were similar across days post-sham procedure and low filament scores did not increase over time.

Second, we determined the weight of filaments that correspond to the break point (when the mouse takes more than 5 minutes to pass its face out of the opening) for both DPI and sham mice for each testing day. As time following pulp exposure increases, the intensity of the filaments where the mice indicate stimulation should stop becomes lower on both sides (Fig.5A,C), such that by day 6 the break point occurs at 0.4g ipsilateral and 0.6g contralateral. In contrast, although there is some change in the number of sham mice that tolerate stimulation with filaments above 0.4g after 7 tests, at least two mice tolerated the entire range of filaments during every test (Fig.5B,D). Taken together this indicates that exposure of one tooth pulp results in a progressive development of mechanical allodynia, which is fully realized on day 4 post-exposure and increases in severity by day 6.

**Figure 5.**
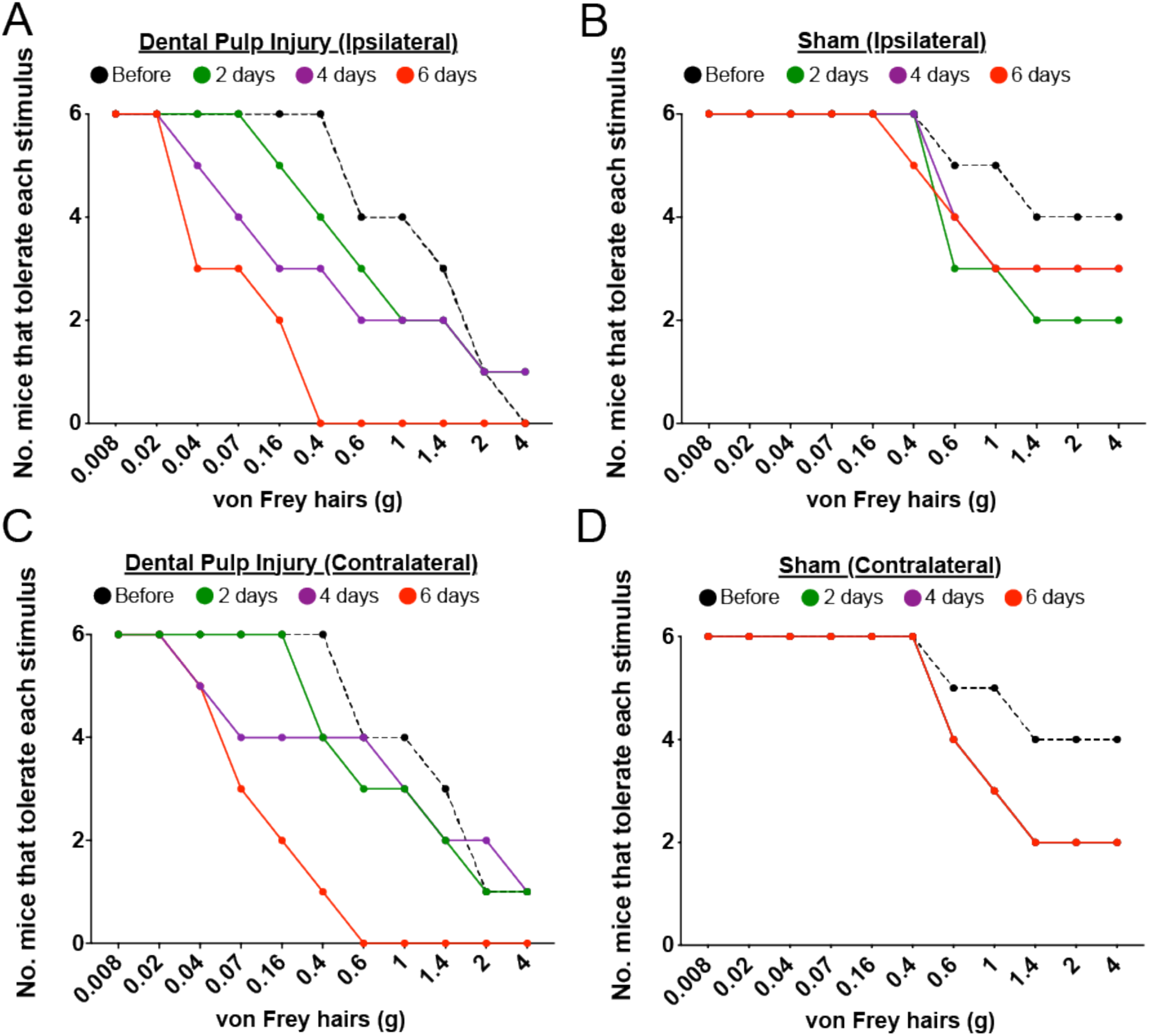
Loss of mouse participation by stimulus intensity (the break point) before and after injury or sham. On the (A) Ipsilateral and (C) contralateral sides, there is a progressive decrease in the number of injured mice willing to tolerate filaments higher than 0.4g, such that by day 6 none of them progressed further than 0.16g (ipsilateral) or 0.4g (contralateral). In contrast, on the (B) ipsilateral and (D) contralateral sides the sham cohort consistently tolerated testing with lower intensity filaments and at least two mice tolerated all the filaments across all testing days.

## Discussion

In this study, we found that unilateral pulp exposure injury in mice to the first maxillary molar resulted in a statistically significant increase in MGS from the first 24 hours onward, which indicates increased spontaneous pain. The pulp was still present at this time, but clearly exposed when examined histologically, supporting that the behavior could capture pain originating from the dental pulp, modeling pulpitis. Surprisingly, mechanical allodynia, as assessed by Von Frey filament testing, progressed more gradually, with initial changes in scores observed only on the contralateral side on day 2 post-injury, significant increases seen on both sides at day 4, and unwillingness to tolerate filaments above 0.6g by day 6 post-injury. This work demonstrates that we have clear easily identifiable behavioral readouts for tooth pain in the mouse. Associated with these behavioral changes, we observed significant increases in transcript levels of Tlr4, Trpv1, and Trpa1, but not Mrgprd, in the ipsilateral TG of injured mice as compared to controls at 24 hours post-injury. Taken together and in support of previous literature, we believe these findings may suggest that Tlr4, Trpv1, and Trpa1, may contribute to early changes resulting in the presentation of spontaneous pain, as indicated by the MGS. However, this does not rule out a role for Mrgprd in the progressive development of mechanical allodynia that seems to worsen around 4 days following pulp-exposure injury. Based on existing literature from other body regions, TRPA1 could serve as a bridge between the early signaling indicated here and a hypothesized later process involving MRGPRD, which we will discuss in further detail below.

Part of our objective in this study was to establish a time course of behavioral changes associated with tooth pulp exposure injury, which is considered by many to be a translationally relevant model for pulpitis^6,31^. To our knowledge, this is the first assessment of MGS following tooth pulp exposure injury, and somewhat surprisingly the first for facial Von Frey in mouse with this model as well. MGS and the rat equivalent RGS are significantly elevated following other types of dental pain, including tooth movement^32^ and mechanical load injury to the temporomandibular joint (TMJ)^33^, but in both of these cases, the elevation in score is transient, likely only corresponding with the presence of acute mechanical load. Our elevation in score does not subside, possibly due to the more invasive nature of the injury and the fact that ongoing inflammation is not being treated. In agreement with this idea, in a chronic model of trigeminal neuropathic pain, significant change in MGS is observed 10 days following the constriction injury^34^. It is possible, however, that there may be site or model specific differences. MGS was only transiently elevated following Complete Freund’s Adjuvant inflammation of the TMJ ^35^, which does not have ongoing infection occurring in the model. In terms of change in the score, our data reflect a similar to slightly greater increase in MGS as compared to tooth movement^32^, and potentially within the lower end of ranges reported for an exogenous-CGRP migraine model^36^ and neuropathic injury of the infraorbital trigeminal nerve^34^. This indicates that the mice are likely in a level of discomfort or pain similar to other experimental pain states. Spontaneous pain is diagnostically associated with irreversible pulpitis, supporting the translational relevance of our findings^29^.

In addition to spontaneous pain, greater than 50% of patients with irreversible pulpitis also have mechanical allodynia with percussion of the tooth, and these patients have higher ratings of spontaneous pain than those without allodynia^29^. Facial Von Frey, an equivalent means of testing mechanical sensitivity in rodents is challenging, but not impossible in the mouse. Mechanical allodynia in the face has been examined in other experimental paradigms, but has not been published following tooth pulp exposure injury. Most studies have used rats as the model animal, and only one of these used the exact model we use here, where the pulp is left exposed and not treated with exogenous substances^31^. Our findings are in agreement with this previous work in rats. Tsuboi and colleagues also observed a reduction in threshold both ipsilateral and contralateral to the injury, first detectable at day 3, which worsened at day 5 and persisted at least 3 weeks later^31^. This period of time around day 3 or day 4 seems to mark a transitional state between the acute inflammatory response and development of pathological pain states often associated with chronic or ongoing pain. We speculate that the early change in MGS may be established by either the same or different mechanisms than those that produce mechanical allodynia later.

To begin to address this question, we examined the mRNA expression of Tlr4, trpv1, Trpa1, and Mrgprd using in situ hybridization at 24 hours following pulp exposure injury. A great deal of attention has been paid to TLR4 as a possible drug target for the treatment of inflammatory pain in various parts of the body^19^, but particularly in pulpitis given its role in recognizing molecular signals of bacterial presence and mechanical injury and upregulation in human pulpitis samples^20^. Furthermore, antagonism of TLR4 is associated with reversal of pain-associated behaviors in two different rat models of pulpitis^7,37^, including mechanical hypersensitivity in lightly anesthetized rats^37^. Our findings of increased Tlr4 in the trigeminal ganglia 24 hours following pulp injury suggest an association between the function of this receptor and at least increased malaise or spontaneous pain associated with increased MGS. We need to directly antagonize TLR4 in the context of pulp-exposure injury to verify causality for increased MGS and determine if early intervention might prevent the delayed presentation of mechanical allodynia. It is possible that TLR4 upregulation begins a cascade of molecular events, as of yet not clearly identified, that establish a change in mechanical sensitivity.

Coinciding with the increase in TLR4 we also observed an increase in Trpv1 mRNA expression at 24 hours post injury, similar to increased protein expression found in rats with pulp exposure or CFA-induced pulpitis models^7,38^. Upregulation of the nociceptive channel TRPV1 has been demonstrated within 24 hours of LPS application to the tooth pulp, but returned to control levels 3 and 5 days later^26^. Furthermore, LPS can directly act on TRPV1+ trigeminal nociceptors via TLR4 signaling^21^. Antagonism of TRPV1 in the CFA model blocks mechanical hypersensitivity in lightly anesthetized rats^38^, suggesting that TRPV1 could be involved in the development of mechanical allodynia in our pulpitis model. However, given the delayed progression of mechanical allodynia reported here, it is likely that other events downstream of the increased TRPV1 expression in the ganglia are also involved in the pulp exposure model.

We also observed an increase in the expression of Trpa1 in the ipsilateral TG at 24 hours post-injury. TRPA1 is also of interest in the pathology of painful pulpitis, but only one other study has examined protein expression following pulp exposure injury in rat molar^39^. They also observed increased expression of TRPA1, but it was not significant until Day 4^39^. Our differing results may be due to species differences, or may reflect a disconnect between the time to peak mRNA levels versus protein levels. Like TRPV1, there is also evidence for an interaction between TLR4 or LPS and TRPA1-related activity. In the DRG, there is evidence that TRPA1 is required for direct nociceptor responses to LPS, even in the absence of TLR4^22^. LPS increases the percentage of trigeminal neurons responding to the TRPA1 agonist acyl-isothiocyanate (AITC) as demonstrated by calcium imaging^23^. TRPA1 has been implicated in the development of mechanical allodynia in the lower body^40^, thus could be involved in the mechanical allodynia reported here.

While we observed increased expression of Tlr4, Trpv1, and Trpa1 24 hours post pulp exposure, we did not observe an increase in Mrgprd, also found in the pulp^25^ and directly implicate in cutaneous mechanical nociception^28,41^. However, this does not completely rule involvement of Mrgprd+ trigeminal neurons in the development of delayed mechanical allodynia. Future studies will evaluate the expression of Mrgprd closer in time to the manifestation of mechanical allodynia around day 3 or 4. It is also possible that TRPA1 could serve as a mechanistic bridge between the early upregulation of TLR4 and TRPV1 and a currently unverified but likely later involvement of Mrgprd following tooth pulp exposure injury. In the DRG recent studies suggest that TRPA1 exists as part of two populations, one that is co-localized with TRPV1 and/or TLR4 in a primarily peptidergic population, which has been the dogma for this channel until very recently, and the other non-peptidergic population containing MRGPRD^42^. Functional studies suggest that at baseline functional TRPA1 protein is actually more frequent in the IB4 positive “non-peptidergic” cell population, which contains MRGPRD, than in the CGRP+ population^42^. The authors suggest that the interaction between TRPA1 and Mrgprd may be broadly important for the development of mechanical allodynia^42^. Very recent evidence suggests that Mrgprd activation by its agonist beta-alanine results in phosphorylation of TRPA1 via Protein Kinase A^43^, but it is not clear if TRPA1 may also influence any aspect of Mrgprd expression or function. The overlap of TRPA1 and Mrgprd in the tooth pulp afferents has not been explored. Alternatively, paracrine signaling in the trigeminal ganglia via gap junction connections with satellite glia^37,38,44^ could allow for recruitment of the non-peptidergic TLR4 negative MRGPRD population by the peptidergic TLR4+/TRPV1 and/or TRPA1+ cell populations to produce the delayed mechanical allodynia we observed.

Although Tlr4 has been heavily examined for its ability to recognize LPS produced by gram-negative bacteria and to influence nociceptor function, it should be cautioned that this is likely not the only potential therapeutic target. Gram negative bacteria are likely to be more involved in the early stages of infection^45^, which may occur before patients make it to the clinic. Bacteria most highly associated with cold and heat sensitive irreversible pulpitis infections are actually gram positive^46^, and may influence pulp nociceptors by different mechanisms than LPS-mediated activation of TLR4 or other TLRs. Other bacterial products^47^ or aspects of inflammation, such as oxidative stress may also recruit both TRP channel expressing nociceptors^48^ and Mrgprd^49^. Clearly, more work is needed on our path to improving the care of endodontic patients.

## Authors Competing Interests Statement

No competing interests to declare.

## Author Contributions

HLR, LPS, JG, BS, CHM, and IAS designed experiments. HLR and LPS carried out experiments, and WF and SM scored florescent intensity of RNA scope images. All authors contributed to the writing and editing of the manuscript.

### Acknowledgements

We thank members of the Abdus-Saboor lab for helpful discussion of this work and comments on this manuscript. This work was supported by startup funds from the University of Pennsylvania to I.A.S. and the National Institutes of Health (NIDCR) R00 grant (DE026807) to I.A.S.

## Methods

### Animals

For these studies we used male and female adult wildtype consisting of a mixed CD1 and C57BL6/J background. Mice were 17-21 weeks old at the time of testing. Mice were maintained in a stardard 12:12 light dark cycle (lights on at 07:00) tested within a time range of 08:30 – 13:00. Mice had access to food and water ad libitum when not being tested. All procedures were approved by the University of Pennsylvania Institutional Animal Care and Use Committee and follow the guidelines established by the National Institutes of Health.

### Dental Pulp Injury

Mice were anesthetized with ketamine/xylazine (i.p. 100 mg/kg and 12.5mg/kg respectively) and positioned under a dissecting microscope and warming pad on their back, with their head supported at an angle, and their mouth propped open with forceps. After trimming the oral whiskers, the upper first maxillary molar was drilled on one side using ¼ round carbide burr until the enamel and dentin layers were breached and the pulp was exposed. This process took about 5 minutes. The enamel is hard and white, the dentin is gray, and when the pulp is visible vasculature and white to pink tissue can be seen in the hole in the enamel under the microscope. Sham animals underwent the same anesthesia, positioning and oral manipulation, but their teeth were not drilled. We provided moist food and monitored body weight following the procedure. Weight loss did not exceed 10%. Mice were either used for behavioral testing on days 1-6 post procedure (n = 6/ treatment group), or were immediately euthanized on day 1 (n =3 injured) to collect tissue. The same set of mice was used for Mouse Grimace Scale and Von Frey testing, performed on alternating days.

### Mouse Grimace Scale

The Mouse Grimace Scale is a scoring system developed in the laboratory of Jeff Mogil to objectively evaluate pain-like facial expressions following experimental procedures^13^, which has been adopted for many trigeminal pain models^33,34,36^, but not yet used to evaluate tooth pain in rodents. Mice (6/treatment group) were acclimated in the chambers at least twice prior to baseline testing, and were in the chamber for 10 minutes before recording began each day. Before the procedure and on days 1, 3, and 5 after injury, we video recorded mice for 10 minutes in clear acrylic chambers (4.3 W × 4.3 H × 11 L cm) on a mesh platform from the small end of the chamber with a camcorder (Sony, HVC) with digital zoom. A 3-way mirror was placed at the back of one end to facilitate assessment of unilateral grooming and to prevent the mouse from viewing the next acclimating mouse. From the 10-minute video, one still image for every second of video was extracted using Video to Picture Converter Software (Hootech). From these ~600 images, 10 were selected that contained a clear view of the animal’s face. All of the 480 selected baseline, sham, and post-pulp exposure images were cropped to show only the face and randomized for scoring in a Power Point file. Scoring was performed blind to day and treatment, as indicated in the original method for 5 action units (orbital tightening, nose bulge, cheek bulge, ear position and whisker change), from 0 (not present) to 2 (very visible), and action units were averaged to arrive at the score for each image^13^. In some cases, the whiskers could not be viewed, so this unit was omitted for the score of that image. Performing the statistical analysis with or without the whisker change action unit did not affect the overall statistical results. For example images of a mouse before and after pulp exposure, see Fig. 2 A,B.

### Mechanical Allodynia Assessment by Von Frey

For these studies we placed the animals in confined chambers with adjustable openings (Fig. 3). The mice were contained chamber about 7cm in all directions, with an opening as wide as 2.5 cm. The animals were acclimated to the chambers once for 30 minutes the day prior to baseline testing. Their natural tendency is to put their face out of the opening when it is wide enough, but they are elevated from the floor, which prevents immediate escape. In this way, we can prompt the animal to present its face for stimulation. We then stimulated twice on either side of the face, alternating between sides, aiming for the region including the vibrissae to the point in front of the eyes. The animal’s response was scored from 0 to 4 based on early work in rats with neuropathic injury ^15,50^ (see Table 1 for score description). We considered “threshold” to be the filament that either produced a score of 3 followed by a response of 2 or more, or the point the animal was no longer willing to pass its face out of the opening after about 5 minutes. The animals were tested on days 2, 4, and 6 post injury with the full filament series (Baseline Tactile Sensory Evaluators, consisting of 11 graded filaments from 0.008g to 4g).

**Table 1.** Score for Responses to Facial Von Frey.

| Score | Response |
| --- | --- |
| 0 | no response |
| 1 | orientation to the stimulus or a slower head turn away from the stimulus |
| 2 | a rapid withdrawal that may or may not be followed by a single facial wipe |
| 3 | attacking or biting the filament or rapid withdrawal followed by 2 facial wipes |
| 4 | a rapid withdrawal with multiple facial wipes |

### Preparation of Tissue for Histology

At 1 day post-injury, sham and naive, mice were deeply anesthetized with ketamine/xylazine and perfused with cold Phosphate Buffered Saline followed by 4% paraformaldehyde through the heart. We removed their trigeminal ganglia and either the remaining cranium or just the portion of the mouth containing the teeth and nerve roots. Trigeminal ganglia and mandible regions were post fixed for up to 4 hours and overnight, respectively. Trigeminal ganglia were placed in 30% sucrose until they sunk (overnight), and then frozen in Neg50 media for cryosectioning (20µm). Teeth were placed in 10% EDTA for approximately two weeks to decalcify, cryoprotected in 30% sucrose, embedded and cryosectioned (20 µm). All tissues were sectioned on a Leica cryostat onto superfrost plus slides, in a series of 16 (TGs) or 10 (teeth). Adjacent series of TG sections were selected for in situ hybridization using the RNAScope system for 2 probes. Four sections per left and right TGs from 3 animals were mounted on one slide. One series from the teeth underwent standard hematoxylin and eosin staining to visualize injury related alterations in the tissue.

### *In Situ* Hybridization using RNAScope

Trigeminal ganglia were prepared using a modified version of the manufacturer’s recommendations for fixed frozen tissues used for fluorescence visualization. Briefly, slides were dehydrated in a graded series of alcohol, peroxidase activity was blocked with hydrogen peroxide, and protease IV was applied to the tissue for 30 minutes at room temperature before undergoing the RNAScope Multiplex Fluorescent v2 assay (ACD). The assay was performed according to the manufacturer’s protocol using two probes. TG sections were assessed for overlap between Tlr4 (channel 1) and either Trpa1, Trpv1, or Mrgprd (channel 2). Channel 1 was visualized using opal dye 520 and channel 2 was visualized with opal dye 570 (1:1500 for both dyes). Tissues were imaged on a Leica SPE TCM using the same laser power and gain settings for all slides. Because we did not know the time course of pain changes in our DPI model, we opted to leave the pulp exposed, rather than applying a dye after exposure and sealing it and the injury site. Thus, we could not be fully certain that the neurons we visualized in the trigeminal ganglion came from the tooth pulp versus other trigeminal tissues. However, our own preliminary studies and others^24^ have shown that maxillary molar labeling with the DiI paste Neurotrace (Invitrogen) results in positive cells in all branches of the trigeminal nerve, therefore we imaged cell clusters observed in both the region where V3 and V2 meet, as well as the region where V1 and V2 meet, resulting in 2 images per section with the 20x objective. All cells with detectable signal were selected for quantification and the signal intensity of mRNA clusters observed within each cell was analyzed by drawing a region of interest around each cell and mean signal intensity in arbitrary units generated by ImageJ software was noted. The dimensions of the region of interest were kept constant throughout the analysis to avoid bias. This process was repeated for each channel including overlay images. The entire quantification was performed by an observer in a manner blinded to mRNA probes and channel assignments.

### Statistical Analysis

Data were assessed for normality using the Shapiro-Wilkes test. Raw Von Frey thresholds were log-transformed to achieve normality so that parametric statistical tests could be used. For Von Frey and Grimace behaviors, two-way ANOVA with repeated measures and between subjects effects were used to determine if there were any significant effects of time, treatment, or a significant interaction, followed by with post-hoc Dunnett’s and Sidak tests where appropriate. For fluorescence intensity, an independent sample t-test was used. We consider p <0.05 significant, and used Graph Pad Prism (v8).

## References

1 Cohen, L. A. et al. Coping with Toothache Pain: A Qualitative Study of Low-Income Persons and Minorities. Journal of Public Health Dentistry 67, 28–35, doi:doi:10.1111/j.1752-7325.2007.00005.x (2007).

2 Erdogan, O., Malek, M., Janal, M. N. & Gibbs, J. L. Sensory testing associates with pain quality descriptors during acute dental pain. European Journal of Pain Jun 26. doi: 10.1002/ejp.1447. [Epub ahead of print], doi:10.1002/ejp.1447 (2019).

3 Nixdorf, D. R. et al. Frequency, impact, and predictors of persistent pain after root canal treatment: a national dental PBRN study. PAIN 157, 159–165, doi:10.1097/j.pain.0000000000000343 (2016).

4 Vena, D. A. et al. Prevalence of Persistent Pain 3 to 5 Years Post Primary Root Canal Therapy and Its Impact on Oral Health–Related Quality of Life: PEARL Network Findings. Journal of Endodontics 40, 1917–1921, doi:https://doi.org/10.1016/j.joen.2014.07.026 (2014).

5 Reda, S. F., Reda, S. M., Thomson, W. M. & Schwendicke, F. Inequality in Utilization of Dental Services: A Systematic Review and Meta-analysis. American Journal of Public Health 108, E1–E7, doi:http://dx.doi.org/10.2105/AJPH.2017.304180 (2018).

6 Gibbs, J. L., Urban, R. & Basbaum, A. I. Paradoxical surrogate markers of dental injury-induced pain in the mouse. PAIN® 154, 1358–1367, doi:https://doi.org/10.1016/j.pain.2013.04.018 (2013).

7 Lin, J.-J. et al. Toll-like receptor 4 signaling in neurons of trigeminal ganglion contributes to nociception induced by acute pulpitis in rats. Scientific Reports 5, 12549, doi:10.1038/srep12549 https://www.nature.com/articles/srep12549#supplementary-information (2015).

8 Shang, L., Xu, T.-L., Li, F., Su, J. & Li, W.-G. Temporal Dynamics of Anxiety Phenotypes in a Dental Pulp Injury Model. Molecular Pain 11, s12990-12015-10040-12993, doi:10.1186/s12990-015-0040-3 (2015).

9 Bradman, M. J. G., Ferrini, F., Salio, C. & Merighi, A. Practical mechanical threshold estimation in rodents using von Frey hairs/Semmes–Weinstein monofilaments: Towards a rational method. Journal of Neuroscience Methods 255, 92–103, doi:https://doi.org/10.1016/j.jneumeth.2015.08.010 (2015).

10 Abdus-Saboor, I. et al. Development of a mouse pain scale using sub-second behavioral mapping and statistical modeling. Cell Reports In press (2019).

11 Lee, C. S. et al. Molecular, cellular, and behavioral changes associated with pathological pain signaling occur after dental pulp injury. Molecular Pain 13, 1744806917715173, doi:10.1177/1744806917715173 (2017).

12 Rodriguez, E. et al. A craniofacial-specific monosynaptic circuit enables heightened affective pain. Nature Neuroscience 20, 1734 (2017).

13 Langford, D. J. et al. Coding of facial expressions of pain in the laboratory mouse. Nature Methods 7, 447+ (2010).

14 Tuttle, A. H. et al. A deep neural network to assess spontaneous pain from mouse facial expressions. Molecular Pain 14, 1744806918763658, doi:10.1177/1744806918763658 (2018).

15 Vos, B., Strassman, A. & Maciewicz, R. Behavioral evidence of trigeminal neuropathic pain following chronic constriction injury to the rat’s infraorbital nerve. The Journal of Neuroscience 14, 2708–2723, doi:10.1523/jneurosci.14-05-02708.1994 (1994).

16 Kartha, S., Zhou, T., Granquist, E. J. & Winkelstein, B. A. Development of a Rat Model of Mechanically Induced Tunable Pain and Associated Temporomandibular Joint Responses. Journal of Oral and Maxillofacial Surgery 74, 54.e51–54.e10, doi:https://doi.org/10.1016/j.joms.2015.09.005 (2016).

17 Dolan, J. C., Lam, D. K., Achdjian, S. H. & Schmidt, B. L. The dolognawmeter: A novel instrument and assay to quantify nociception in rodent models of orofacial pain. Journal of Neuroscience Methods 187, 207–215, doi:https://doi.org/10.1016/j.jneumeth.2010.01.012 (2010).

18 Hall, B. E. et al. Conditional TNF-α Overexpression in the Tooth and Alveolar Bone Results in Painful Pulpitis and Osteitis. Journal of Dental Research 95, 188–195, doi:10.1177/0022034515612022 (2015).

19 Bruno, K. et al. Targeting toll-like receptor-4 (TLR4)-an emerging therapeutic target for persistent pain states. Pain 159, 1908–1915, doi:10.1097/j.pain.0000000000001306 (2018).

20 Wadachi, R. & Hargreaves, K. M. Trigeminal Nociceptors Express TLR-4 and CD14: a Mechanism for Pain due to Infection. Journal of Dental Research 85, 49–53, doi:10.1177/154405910608500108 (2006).

21 Diogenes, A., Ferraz, C. C. R., Akopian, A. N., Henry, M. A. & Hargreaves, K. M. LPS Sensitizes TRPV1 via Activation of TLR4 in Trigeminal Sensory Neurons. Journal of Dental Research 90, 759–764, doi:10.1177/0022034511400225 (2011).

22 Meseguer, V. et al. TRPA1 channels mediate acute neurogenic inflammation and pain produced by bacterial endotoxins. Nature Communications 5, 3125, doi:10.1038/ncomms4125 https://www.nature.com/articles/ncomms4125#supplementary-information (2014).

23 Michot, B., Casey, S., Lee, C. & Gibbs, J. (135) - LPS-induced neuronal activation and TRPA1 sensitization in trigeminal sensory neurons is dependent to TLR4 receptor. The Journal of Pain 19, S10–S11, doi:https://doi.org/10.1016/j.jpain.2017.12.049 (2018).

24 Kadala, A. et al. Fluorescent Labeling and 2-Photon Imaging of Mouse Tooth Pulp Nociceptors. Journal of Dental Research 97, 460–466, doi:10.1177/0022034517740577 (2018).

25 Chung, M.-K., Jue, S. S. & Dong, X. Projection of Non-peptidergic Afferents to Mouse Tooth Pulp. Journal of Dental Research 91, 777–782, doi:10.1177/0022034512450298 (2012).

26 Chung, M.-K., Lee, J., Duraes, G. & Ro, J. Y. Lipopolysaccharide-induced Pulpitis Up-regulates TRPV1 in Trigeminal Ganglia. Journal of Dental Research 90, 1103–1107, doi:10.1177/0022034511413284 (2011).

27 Michot, B., Lee, C. S. & Gibbs, J. L. TRPM8 and TRPA1 do not contribute to dental pulp sensitivity to cold. Scientific Reports 8, 13198, doi:10.1038/s41598-018-31487-2 (2018).

28 Cavanaugh, D. J. et al. Distinct Subsets of Unmyelinated Primary Sensory Fibers Mediate Behavioral Responses to Noxious Thermal and Mechanical Stimuli. Proceedings of the National Academy of Sciences of the United States of America 106, 9075–9080 (2009).

29 Owatz, C. B. et al. The Incidence of Mechanical Allodynia in Patients With Irreversible Pulpitis. Journal of Endodontics 33, 552–556, doi:https://doi.org/10.1016/j.joen.2007.01.023 (2007).

30 Renton, T. & Wilson, N. H. Understanding and managing dental and orofacial pain in general practice. British Journal of General Practice 66, 236–237, doi:10.3399/bjgp16X684901 (2016).

31 Tsuboi, Y. et al. Modulation of astroglial glutamine synthetase activity affects nociceptive behaviour and central sensitization of medullary dorsal horn nociceptive neurons in a rat model of chronic pulpitis. European Journal of Neuroscience 34, 292–302, doi:10.1111/j.1460-9568.2011.07747.x (2011).

32 Zhu, Y. et al. Effect of static magnetic field on pain level and expression of P2X3 receptors in the trigeminal ganglion in mice following experimental tooth movement. Bioelectromagnetics 38, 22–30, doi:10.1002/bem.22009 (2017).

33 Sperry, M. M., Yu, Y.-H., Welch, R. L., Granquist, E. J. & Winkelstein, B. A. Grading facial expression is a sensitive means to detect grimace differences in orofacial pain in a rat model. Scientific Reports 8, 13894, doi:10.1038/s41598-018-32297-2 (2018).

34 Akintola, T. et al. The grimace scale reliably assesses chronic pain in a rodent model of trigeminal neuropathic pain. Neurobiology of Pain 2, 13–17, doi:https://doi.org/10.1016/j.ynpai.2017.10.001 (2017).

35 Bai, Q. et al. TNFα in the Trigeminal Nociceptive System Is Critical for Temporomandibular Joint Pain. Molecular Neurobiology 56, 278–291, doi:10.1007/s12035-018-1076-y (2019).

36 Rea, B. J. a. et al. Peripherally administered calcitonin gene-related peptide induces spontaneous pain in mice: implications for migraine. Pain 159, 2306–2317 (2018).

37 Ohara, K. et al. Toll-like receptor 4 signaling in trigeminal ganglion neurons contributes tongue-referred pain associated with tooth pulp inflammation. Journal of Neuroinflammation 10, 139, doi:10.1186/1742-2094-10-139 (2013).

38 Watase, T. et al. Involvement of transient receptor potential vanilloid 1 channel expression in orofacial cutaneous hypersensitivity following tooth pulp inflammation. Journal of Oral Science advpub, doi:10.2334/josnusd.16-0854 (2018).

39 Haas, E. T., Rowland, K. & Gautam, M. Tooth injury increases expression of the cold sensitive TRP channel TRPA1 in trigeminal neurons. Archives of Oral Biology 56, 1604–1609, doi:https://doi.org/10.1016/j.archoralbio.2011.06.014 (2011).

40 Lennertz, R. C., Kossyreva, E. A., Smith, A. K. & Stucky, C. L. TRPA1 Mediates Mechanical Sensitization in Nociceptors during Inflammation. PLoS ONE 7, e43597 (2012).

41 Zylka, M. J., Rice, F. L. & Anderson, D. J. Topographically Distinct Epidermal Nociceptive Circuits Revealed by Axonal Tracers Targeted to Mrgprd. Neuron 45, 17–25, doi:https://doi.org/10.1016/j.neuron.2004.12.015 (2005).

42 Barabas, M. E., Kossyreva, E. A. & Stucky, C. L. TRPA1 Is Functionally Expressed Primarily by IB4-Binding, Non-Peptidergic Mouse and Rat Sensory Neurons. PLoS ONE 7, e47988 (2012).

43 Wang, C. et al. Facilitation of MrgprD by TRP-A1 promotes neuropathic pain. The FASEB Journal 33, 1360–1373, doi:10.1096/fj.201800615RR (2019).

44 Komiya, H. et al. Connexin 43 expression in satellite glial cells contributes to ectopic tooth-pulp pain. Journal of Oral Science 60, 493–499, doi:10.2334/josnusd.17-0452 (2018).

45 Jang, J.-H. et al. An Overview of Pathogen Recognition Receptors for Innate Immunity in Dental Pulp. Mediators of Inflammation 2015, 12, doi:10.1155/2015/794143 (2015).

46 Zheng, J. et al. Microbiome of Deep Dentinal Caries from Reversible Pulpitis to Irreversible Pulpitis. Journal of Endodontics 45, 302-309.e301, doi:https://doi.org/10.1016/j.joen.2018.11.017 (2019).

47 Chiu, I. M. et al. Bacteria activate sensory neurons that modulate pain and inflammation. Nature 501, 52+ (2013).

48 Hargreaves, K. M. & Ruparel, S. Role of Oxidized Lipids and TRP Channels in Orofacial Pain and Inflammation. Journal of Dental Research 95, 1117–1123, doi:10.1177/0022034516653751 (2016).

49 Bautzova, T. et al. 5-oxoETE triggers nociception in constipation-predominant irritable bowel syndrome through MAS-related G protein–coupled receptor D. Science Signaling 11, eaal2171, doi:10.1126/scisignal.aal2171 (2018).

50 Deseure, K., Koek, W., Adriaensen, H. & Colpaert, F. C. Continuous Administration of the 5-Hydroxytryptamine<sub>1A</sub> Agonist (3-Chloro-4-fluoro-phenyl)-[4-fluoro-4-{[(5-methyl-pyridin-2-ylmethyl) -amino]-methyl}piperidin-1-yl]-methadone (F 13640) Attenuates Allodynia-Like Behavior in a Rat Model of Trigeminal Neuropathic Pain. Journal of Pharmacology and Experimental Therapeutics 306, 505–514, doi:10.1124/jpet.103.050286 (2003).

